# Enhanced Cancer Subtyping via Pan-Transcriptomics Data Fusion, Monte-Carlo Consensus Clustering, and Auto Classifier Creation

**DOI:** 10.1101/2019.12.16.870188

**Authors:** Kristofer Linton-Reid, Harry Clifford, Joe Sneath Thompson

## Abstract

Subtyping of tumor transcriptome expression profiles is a routine method used to distinguish tumor heterogeneity. Unsupervised clustering techniques are often combined with survival analysis to decipher the relationship between genes and the survival times of patients. However, the reproducibility of these subtyping based studies is poor. There are multiple reports which have conflicting subtype and gene-survival time relationship results. In this study, we introduce the issues underlying the lack of reproducibility in transcriptomic subtyping studies. This problem arises from the routine analysis of small cohorts (< 100 individuals) and use of biased traditional consensus clustering techniques. Our approach carefully combines multiple RNA-sequencing and microarray datasets, followed by subtyping via Monte-Carlo Consensus Clustering and creation of deep subtyping classifiers. This paper demonstrates an improved subtyping methodology by investigating pancreatic ductal adenocarcinoma. Importantly, our methodology identifies six biologically novel pancreatic ductal adenocarcinoma subtypes. Our approach also enables a degree of reproducibility, via our pancreatic ductal adenocarcinoma classifier PDACNet, which classical subtyping studies have failed to establish.

## 1 INTRODUCTION

Subtyping of tumoral transcriptomic expression data is a key method in cancer informatics. These studies involve division of heterogeneous tumour populations into clinically and biologically distinct subtypes. Tumour tissues, from the same location, that appear morphological similar may have significantly different molecular features. This may attribute to variable responses to therapy and clinical outcomes. Once molecular subtypes of a cancer are defined, they can guide the use of therapies and treatment options, within trials and clinically.

Tumoral subtypes are usually distinguished by employing unsupervised consensus clustering techniques on expression datasets. After clustering, these subtypes are typically biologically validated via Kaplan-Meier survival analysis. This infers how expression changes and analogous subtypes affect the treatments and progression of cancer.

There are several gene expression based subtyping studies that cover a range of cancer types. However, a bottleneck here is the inconsistency between studies. Different cohorts and clustering techniques can produce very different subtyping results. Reports on the clustering of epithelial ovarian cancer have distinguished 4 to 6 subtypes [2,12,29,32]. Similarly, colorectal cancer has been classified into 3 to 6 subtypes [10]. Pancreatic cancer has been considered to comprise of 2 to 6 subtypes [[1,3,5,21,34,38]].

With the multiple inconsistencies in consideration, the current framework of subtyping needs developing further. There are two main weaknesses in the majority of subtyping studies. These stem from the use of small cohorts (<100 samples) and the use of unsupervised consensus clustering techniques, usually non-negative matrix factorisation (NMF), to distinguish subtypes.

The issue with using small cohorts is that each subtype may only be made up of a few stable samples, meaning only a few samples ‘fit’ into the cluster. Sample stability is indicated by silhouette widths, these values are a measurement of how similar a sample is to its own cluster compared to other clusters. It is clear clusters made up of only a few samples with a silhouette width >0.5 (out of 1) would not represent a population accurately. It is unlikely that analysing expression data of small cohorts will accurately distinguish rarer subtypes. Dieci et al’s study on breast cancer [4] demonstrated that rare subtypes of breast cancer exist with distinct molecular profiles and responses to treatment. It is clear that more comprehensive subtyping studies must be conducted in order to improve precision medicine treatment decisions.

The main problem with traditional consensus clustering techniques is that they are prone to discovering false positives. In other words, they may indicate the incorrect number of clusters (K). The false positive issue rises from three main weaknesses. The first two weaknesses arise from issues of the clustering techniques. Traditional consensus clustering has an inability to reject the null hypothesis that K=1; and is subject to the inherent stability bias, defined as the tendency of cluster stability to increase as the number of clusters increases (and as the number of samples per cluster is reduced). The third issue originates from the RNA sequencing data itself. This sequencing data has a negative binomial distribution. Pacheco et al. [22] distinguished that negatively binomial distributed data can be classified into varying clusters of various sample sizes, and this dramatically alters performance on cluster-level t-tests. The traditional clustering techniques will fail to account for the over-dispersion of RNA-seq expression data. For example, the popular NMF clustering technique has been reported to produce mixed results that deviate depending upon the starting point [8].

Stemming from the traditional subtyping study design issues, there are reports on inconsistencies existing between studies. There is a challenge in reproducing clusters from the same datasets using different techniques. John et al., [40] displayed reproducibility issues in 5 cancer NMF based subtyping studies, using their novel clustering technique Monte Carlo Consensus Clustering (M3C).

Despite these clear reproducibility issues, there have been no attempts to directly address them. There are three key improvements that can be implemented to remove the bottleneck of the current gene expression subtyping pipeline. The first is to increase the number of patients involved in each study, which should include incorporating multiple international cohorts and increasing the number of samples per cluster. The second is to perform a more robust subtyping method, which rejects the null hypothesis that K=1, and accounts for the inherent stability bias. The third is to build classifiers based upon the subtyping annotations, which to an extent would allow subtyping of ‘new’ samples being added to a cohort post clustering.

In this study we employed our enhanced subtyping methodology on a pancreatic cancer dataset. Specifically, we focused on the pancreatic ductal adenocarcinomas (PDAC) subtype expression profiles, derived from tumor biopsies.

We created the largest open source transcriptomic pancreatic ductal adenocarcinoma dataset to date (1013 patients) by combining open source microarray and RNA-Seq datasets (Table 1). This addresses one of the main challenges when subtyping small cohorts, sample bias. This is a clear problem with PDAC cohorts as 80% of individuals are diagnosed at late stages [17]. Small cohorts typically capture late stage PDAC, and as previously mentioned cannot form accurate and stable clusters.

To distinguish subtypes, we employed John et al’s novel M3C algorithm [40]. This technique is an improvement from traditional clustering techniques as it both allows rejection of the null hypothesis that there is only one subtype and removes the inherent stability bias.

As well as removing the small cohort issue and clustering problems, we performed the standard subtype clinical and biological validation. This was addressed via a Kaplan-Meier survival analysis and differential expression (DE) analysis.

The last step was our novel use of subtype annotations to build a deep learning classifier PDACNet. The creation of tumoral subtype classifiers is ideal for reproducing subtyping results on other tumoral cohorts.

## 2 METHODS

We developed the largest open source transcriptomic pancreatic ductal adenocarcinoma dataset (section 2.1) and created a novel pipeline for gene expression subtyping (section 2.2).

### 2.1 PDAC Dataset Creation

To create this large data set of PDAC expression values, we gathered data from a variety of repositories. These repositories included the International Cancer Genome Consortium (ICGC, www.icgc.org), the Cancer Genome Atlas (TCGA, http://cancergenome.nih.gov/), and Pubmed’s Gene Expression Omnibus (GEO, http://www.ncbi.nlm.nih.gov/geo/). This dataset was created by combining 11 publicly available PDAC Microarray datasets, and 3 RNA-sequencing datasets derived from solid tumor biopsies (Table 1). The 11 publicly available Microarray datasets were merged by matching gene IDs and then batch corrected via ComBat [15], resulting in a dataset of 710 PDAC samples. The same technique was used for the 3 RNA-Sequencing datasets, forming a dataset of 303 samples. The RNA-Sequencing dataset was then normalized to the Microarray dataset, using feature specific quantile normalization [6,7]. Principal Component Analysis (PCA) score plots and Scree Plots, for Microarray and RNA data set integration are in Appendix Figures 1 and 2.

**Table 1.** Sources of data. All data used in this study.

| Reference | Number of Samples | Source |
| --- | --- | --- |
| **Microarray** |  |  |
| Collinson et al., [3] | 27 | Collinson |
| Ishikawa et al., [13] | 49 | GSE1542 |
| Liadaki et al., [19] | 39 | GSE1571 |
| Sandhu et al., [25,26] | 49 | GSE60980 |
| Janky et al., [14] | 118 | GSE62165 |
| Yang et al., [37] | 69 | GSE62452 |
| Moffit et al., [21] | 145 | GSE71729 |
| Guttman et al., [11] | 80 | GSE8591 |
| Lunardi., [20] | 53 | GSE55643 |
| Zhang et al., | 45 | GSE28735 |
| Wood et al., [28] | 36 | GSE1615 |
| **RNA** |  |  |
| TCGA | 183 | tcga (April 2019) |
| Bailey et al., [1] | 69 | Bailey et al |
| Kirby et al., [15] | 51 | GSE79670 |

**Figure 1.**
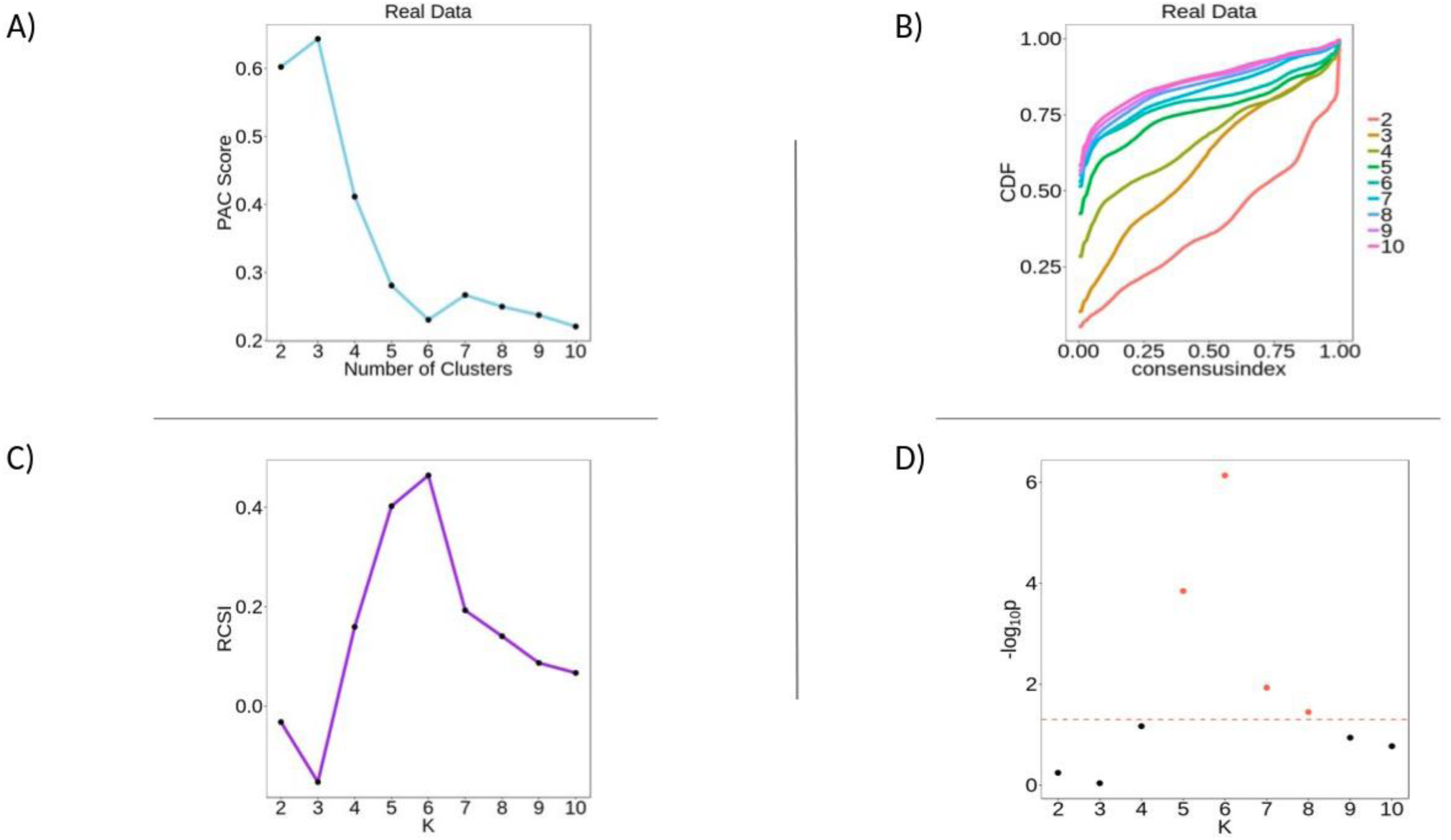
M3C Identification of 6 PDAC Clusters. A) PAC scores display a sharp downward spike at K=6. B) CDF plot shows flattening of the consensus index curve as K increases. C) RSCI shows a sharp upward spike at K=6. D) Plot of the P-values over K=0:10.

**Figure 2.**
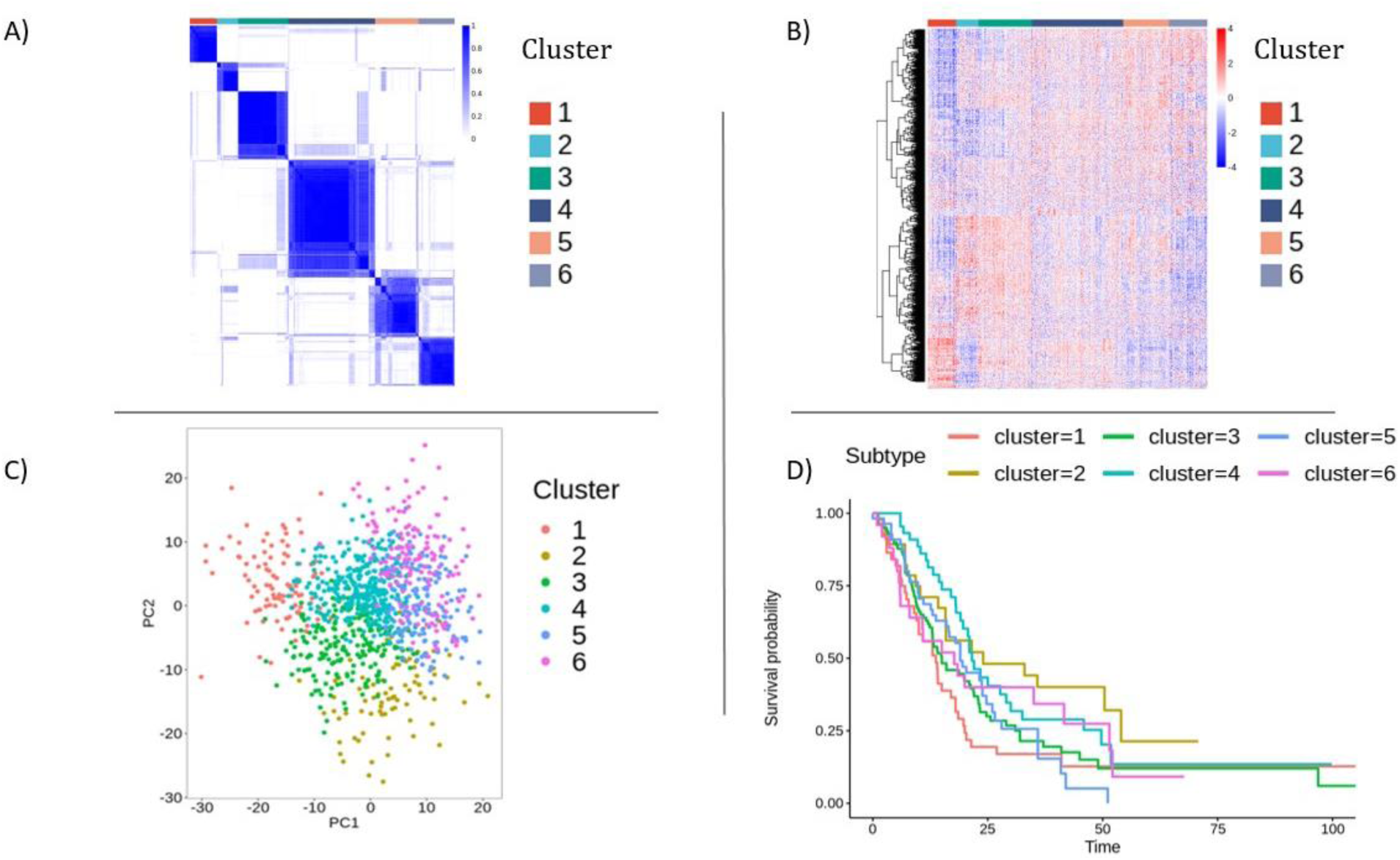
Clinical Validation of PDAC subtypes. A) Consensus matrix for the 6 distinct subtypes. B) Heatmap of the 2000 most variable genes, displaying distinct expression profiles for each subtype. C) PCA score plot (Components 1 & 2) of the 6 subtypes, subtypes 5 and 6 have the most overlap. D) Kaplan-Meier curves showing differing survival trends for the identified subtypes P-value= 0.063.

### 2.2 Subtyping Pipeline

The subtyping approach encompasses 4 key components. 1) Clustering with M3C, 2) Biological validation via Kaplan-Meier survival analysis, 3) Differential Expression and Gene Set Enrichment Analysis, 4) Subtype Classifier Creation.

#### 2.2.1 Clustering

Detailed by John et al., (2018) [40], the improvements of the M3C algorithm are all based upon the use of Monte-Carlo simulations. These are used to create a reference matrix. In brief, the simulations run multiple PCAs, whereby a random scores matrix of the nth simulation is generated, and the eigenvector matrix is calculated. The simulated PCA random scores matrix and eigenvector matrix is then multiplied. These calculations are then repeated for the defined number of simulations. This results in a reference matrix that captures the associations between samples, but without the ‘real’ clusters.

This reference data set is then passed into the consensus clustering algorithm, as well as the original input data, and the two results are compared. The final output of M3C is essentially an improved consensus clustering result, that accounts for the null distribution (reference matrix).

In short, clustering finds samples of similarity and places them into groups without any similarity overlap between groups. The consensus clustering combines resampling with clustering, and integrates all the results from the various runs, in one final cluster result.

There are multiple different clustering and consensus clustering algorithms. The M3C algorithm calculates a novel metric, the Relative Cluster Stability Index (RCSI), obtained from real and simulated proportions of ambiguously clustered (PAC) scores. The RSCI provides a more accurate representation of stability across the distribution of K, as the inherent stability bias is accounted for (See Figures 1A and 1C). The p-values for each k value are derived by comparing the simulated reference consensus clustering results and the real results. If these p-values are significant, then the null hypothesis that K=1 (95% confidence interval) is rejected. In other words, unlike the typical consensus clustering methods, the M3C technique allows rejection of the hypothesis that there are no subtypes. In this study, M3C’s default clustering loop was employed, partitioning around medoids (PAM) with Euclidean distance, with 100 iterations.

#### 2.2.2 Biological validation via Kaplan-Meier survival analysis

Out of the expression cohort of 1013 individuals there were 303 patients with corresponding clinical data available. The Kaplan– Meier technique was employed, which calculates median survival (the shortest time at which the survival probability drops to 50% or lower) as described by Goel et al. [9], and the difference was tested using the log-rank test. P-values of less than 0.05 were considered statistically significant. This was conducted using the survival R package and plotted with the survminer package [31].

#### 2.2.3 Differential Expression

The differential expression analysis was conducted, comparing one subtype to all other subtypes. The limma package ‘lmFit’ [24] function was used to calculate the fold changes and p-values of all genes. These are calculated by employing multiple linear models, on log-transformed expression data, for each comparison. To conduct these comparisons, multiple least squares regressions were employed. These can compare each subtype versus all other subtypes.

#### 2.2.4 Subtype Classifier

A deep learning subtype classifier was created to enable the subtype identification of ‘novel’ PDAC expression samples. This classifier was built by employing the R version of the H2O package ‘autoML’ function on the 500 most variable genes of the 1013 sample dataset. This dataset was randomly partitioned by 1/7th into training and test sets. The ‘autoML’ function is used to create multiple models fully automatically. This technique was set to generate 100 classification models, which included random forest models, XGBoost, deep learning, and stack ensemble models. The model with the lowest mean per class error rate (lowest average subtype identification error), a random grid search deep learning model, was selected as ‘PDACNet’.

In brief, the deep learning subtype classifier is comprised of one hidden layer of 500 neurons that learns through the activation method known as rectifier with dropout (input dropout ratio = 0.15, output dropout ratio = 0.4), with hyperparameters optimised with random grid search (epochs = 46.44). For further details of the deep learning model, see PDACNet file available at the GitHub link in the Data Availability section.

## 3 RESULTS

The results derived from analysis of the large PDAC cohort of 1013 patients. Principle Components Analyses Score and Scree plots (Appendix Figures 1 & 2). In Appendix Figures 1 and 2 there is an increase in variability in components 1 and 2, corresponding to the RNA sequencing and microarray data sets.

### 3.1 M3C Identifies Six Clusters of PDAC

We employed M3C on the dataset of 1013 patients, which resulted in 6 well-defined clusters.

The PAC score plot (Figure 1A) displays a sharp spike at K=6, suggesting this is the optimal number of clusters. However, as with the CDF plot, the inherent stability bias can be seen here as it naturally tends towards lower values as K increases. Importantly, the traditional PAC score and Cophenetic coefficient plots cannot reject the null hypothesis K=1.

The Cumulative Distribution Function (CDF) plot (Figure 1B), is a plot of the consensus matrices. The optimal value of K is the one with the flattest curve. Of note, in the CDF plot there is a noticeable bias of traditional consensus clustering techniques. This bias is based on the increase in stability as K increases for any given dataset.

The M3C approach outputs a Relative Cluster Stability Index (RCSI), which accounts for both PAC scores and simulated reference PAC scores. The RSCI is an improved metric for determining optimal K as it eliminates the inherent stability bias. The RSCI peaked at K=6, confirming there are 6 clusters (Figure 1C).

As part of M3C analysis a beta p-value distribution is calculated. This is used to reject the null hypothesis that K=1. If none of the P-values are significant over a logical distribution of K values, the null hypothesis is accepted. In this case, ranks 5:8 were of significance. The value of K=6 was accepted. A Pearson’s correlation was employed to check that these 6 clusters were not unfairly biased to the initial data sets, we also identified which proportions of the 14 datasets/batches the 6 subtypes comprise of. Appendix: Figure 3 and Table 3). Out of the 1013, 1002 samples fitted into the 6 clusters.

A direct comparison between the traditional NMF clustering technique and the M3C clustering technique was also employed on the pancreatic cancer cohorts from Collisson et al., [3] and Bailey et al., [1] (Appendix Figures 4 & 5). The PAC scores plots indicate that there are 3 and 4 clusters respectively, matching the results documents in their respective papers. However, for both studies, the RSCI metric would indicate that there are 4 and 8 clusters. Furthermore, the corresponding p-value distributions, with no significant values (>alpha=0.05), suggest that the clusters identified here are not particularly strong, hence the issues with reproducibility of these clusters.

### 3.2 Clinical Validation of PDAC subtypes

From the merged dataset of 1013 samples, there are 303 samples with corresponding clinical data. The subtype labels and associated overall survival information were used to perform survival analysis. Figure 2A shows the consensus matrix of the 6 distinct subtypes. Figure 2B, a heatmap of the 2000 most variable genes, displays distinct expression profiles for each subtype. Figure 2C is PCA score plot (Components 1 &2), of the six subtypes, whereby subtypes 5 and 6 have the most overlap. Figure 2D displays the Kaplan-Meier curves. While this was not significant, P-value= 0.063, there is a clear trend that subtype/cluster 5 has the poorest survival outcome, and cluster 3 has the highest number of individuals at risk at time 0.

### 3.3 Functional Identification of PDA subtypes: Differential Expression and Literature Comparisons

Differential expression analysis revealed distinct expression profiles for each subtype. Appendix Table 1 displays the 5 genes with the most significantly altered expression levels in each subtype. Full list of subtype specific genes is available at the GitHub link in the Data Availability section, as ‘PDAC_Subtype_Specific_Genes.csv’).

### 3.4 Subtype Classifier: Selection

In this study, 100 machine learning-based classifiers were created. Importantly the goal of this component was to create a well-performing subtype classifier without the need for expert parameter tuning knowledge. H2O’s autoML function is one of the simplest ways to do this, the minimal required inputs of the function are; an input training and test set, with outcome variable defined. Out of the 100 classifiers, the classifier with the lowest mean per class error was selected to be further analysed (PDACNet). Notably the logarithmic loss result is greater than 1, this combined with the low mean per class error indicates that the model is highly confident about an incorrect subtype classification. Table 2 displays the results of the top 5 classifiers, ordered by the mean per class error. Appendix Table 2 displays the prediction errors when the PDACNet was applied on the test set, the partitioned 1/7th of the 1002 samples that fitted into one of the 6 clustes. For full information regarding parameters of the deep learning model that has the lowest mean per class error, see the PDACNet file available at the GitHub link in the Data Availability section. The important takeaway here is that subtype classifiers can enable the subtype classification on ‘novel’ PDAC expression samples/cohorts that are outside of this study. This enables all the improvements from the information captured by the large cohort and application of M3C to be transferred over when trying to distinguish the subtypes of only a few samples. Applying a subtyping technique to a few samples or a small cohort would provide poorly clustered results.

**Table 2.** The top 5 Subtype Classifiers. Ordered by the mean per class error. The top model is ‘DeepLearning_Drid_Search_Model_1’

| Model id | Mean per Class error | Log Loss | Root mean standard error |
| --- | --- | --- | --- |
| DeepLearning_Grid_Search_Model_1 | 0.140 | 1.09 | 0.385 |
| DeepLearning_Grid_Search_Model_8 | 0.146 | 1.52 | 0.380 |
| Stacked_Ensemble_Best_Of_Family | 0.157 | 0.452 | 0.355 |
| Stacked_Ensemble_All_Models | 0.161 | 0.526 | 0.404 |
| DeepLearning_Grid_Search_Model_8 | 0.173186 | 1.03 | 0.390 |

## 4 DISCUSSION

This work describes an improved workflow for gene expression subtyping studies. This workflow includes rapid creation of expression datasets from both RNA-Sequencing and Microarray datasets. These large datasets can then be subtyped using M3C, which as previously mentioned removes the inherent stability bias and can disprove the null hypothesis that K=1. Biological and clinical validation can be employed by a) Distinguishing differentially expressed genes in each subtype, b) Distinguishing distinct survival times via survival-analysis. Lastly, classifier creation with subtype annotations is a novel way to mimic an unsupervised subtyping study on distinct datasets.

### 4.1 Alternative Methods to Increase Cohort Size

This study focused on a dataset of 1013 PDAC patients, whereby expression data was derived cancer tissue. However, there are multiple studies which have included expression values from tissue and other sources e.g. tumour derived cell lines. While this may increase the statistical power, and robustness, there are reports which indicate not all tumoral cell lines accurately represent solid tumours. This issue was first distinguished in a lung cancer subtyping study, whereby out of the 11 cell lines representing lung adenocarcinomas, none of them formed clusters with tumoral adenocarcinomas [33].

Along the lines of including samples from other sources, other studies have focused on specific cancers, and not a specific type of histopathology. For example, Bailey et.al., [1] performed a subtyping study on six histopathological types of pancreatic cancer. This included: pancreatic adenocarcinomas, adenosquamus, acinar cell carcinoma, intraductal papillary mucinous neoplasm, and four other rare pancreatic tumours. However, as indicated by the silhouettes the sample with the highest stability in their cluster 1 was Acinar cell carcinoma, and in cluster 3 was an adenosquamous carcinoma. Similarly, all 11 intraductal papillary neoplasms were unstable with silhouette widths < 0.1. It appears to identify subtypes accurately on one histopathological type at a time. Intentionally or unintentionally subtyping (through pathological identification errors) different histopathological tumours may also be contributing to the challenge of reproducing subtyping results.

### 4.2 Molecular Discussion of Subtypes

Class 1 included AOX1, FLRT2, SLC1A2, GAMT and KCNJ5. AOX1 is known for being the precursor of a xenobiotic metabolising enzyme, and Sigruener et al., [28] has previously proven downregulation in pancreatic cancer, with its cellular functions revolving around lipid efflux and phagocytosis in hepatocytes. Expression of KCNJ5 has also previously been shown to be downregulated in PDAC compared to controls [30]. This gene is known of the potassium inwardly rectifying channel family.

Interestingly, classes 2 and 3 appear to have the same 5 genes that have the most significantly altered expressions. In class 2 they are upregulated and in class 3 they are downregulated. GAMT which has altered expression in classes 1, 2, and 3 is known to be involved in p53-dependent apoptosis and is particularly important for cell apoptosis and survival under nutrient deficient conditions [39]. P2RX1 gene is the precursor of an ATP-gated nonselective ion channel [16]. NUCB2 has been reported to inhibit apoptosis of pancreatic cells.

In class 4 EPB41L4B is the most significant gene, this plays a role in the proliferation of epithelial cells and matches reports of its downregulation in PDAC [35]. In class 5, the most differentially expressed gene AP1M2 has also been previously reported to be differentially expressed in PDAC stromal tissue [23].

A study by Zhao et al. [38] also identified 6 PDAC subtypes. Interestingly their gene expression analysis was not too dissimilar to this study. They displayed a subtype with carbohydrate metabolism gene expression alterations, such as ALDOB, CA2, NPC1L1 and PGC. Another subtype identified by Zhao et al, appeared to have more alterations in Cell proliferation and epithelium-associated genes, such as CCNB2, CDKN2A, SFN, UBE2C, SPRR3, DHRS9 and CRABP2. GREM1, MFAP5, COL12A1, COL10A1, COL8A1 and other collagen or ECM-related genes. They also identified a subtype with Immune related genes alterations such as CCL, CCR7 and CD gene families. Neuroendocrine-associated genes such as PAX6, IAPP, G6PC2, ABCC8 and ZBTB16 are highly expressed in another subtype. Lastly, Zhao et al. identified a subtype with multiple gene alteration differences involved in lipid and protein metabolism CLPS, PLA2G1B, CEL, ALB, CPA1, CPB1, CTRL, SLC3A1, PRSS3 and ANPEP. Whilst Zhao’s study was rather comprehensive with their gene set enrichment analysis, the original differential expression was based upon inherent differences between subtypes, an alternative approach would compare subtypes to healthy controls. In this case comparing gene expression profiles here is a challenge and would require full data sets of expressed genes to be available.

Overall, differential expression genes are to be expected all have either a function relating to increased cancer likelihood or be attributed dysfunctional pancreatic cells (e.g GAMT and its relationship with p53, and P2RX1 and its ion channel dysfunction).

### 4.3 Overview

To enable the use of our classification on new cohorts we made a subtyping classifier, based upon a large multi-centred dataset. Unlike traditional classification studies, whereby the typical aim is to discover a series of predictive biomarkers, the goal of the classifier creation was to: a) facilitate our subtyping of PDAC on novel cohorts, b) Develop a pipeline that could be easily applied on a different expression cohort, of a different cancer, in a matter of hours (hence no manual fine tuning of parameters in this case). Despite our creation of the multi-centred dataset and M3C results, RSCI and p-value for 6 clusters, it is still possible that the clusters do not fit all PDAC individuals. Hence the classifier could possibly misidentify rare PAC types. Ideally with the widening access of sequencing technologies, our study will be repeated with a larger cohort resembling the PDAC population more accurately.

As well as advances in the amount of expression data available, there will likely be advances in available associated omics data. Expression data does not capture the effects tumoral mutations have on downstream functionality. In other words, a gene could be highly expressed but downstream non-functional. There are multiple data sources which could combine with mRNA-expression data to improve subtyping information capacity. Kuijer et al., [18] suggested that somatic mutation-based subtyping can provide novel insights. Typically, clinical trials use single gene mutations as guidelines. Unique patterns of somatic mutation information combined with mRNA expression subtyping may provide an extra layer to identify individuals who will respond optimally (and individuals who will not respond) to certain treatments. The iCluster algorithm was built to identify novel subtypes by integrating DNA copy number changes and gene expression, which has distinguished novel subtypes in lung and breast cancer [27]. Notably iCluster has been applied to a cohort of 363 pancreatic hepatocellular carcionmas[36], identifying 3 subtypes. However, iCluster is based upon K-means clustering, which is also subject to the inherent stability bias. Perhaps, the addition of a reference matrix and Monte-carlo simulations to K-means clustering could be an integral step in this case.

## 5 CONCLUSION

In conclusion, we have developed a novel approach to subtyping expression data. This includes the generation of large cohorts, use of M3C, DE analysis, and classifier creation. Importantly, our robust distinction of six PDAC subtypes has set a benchmark for future PDAC subtyping studies. This could be a foundation to discovering novel PDAC personalized therapies and improving survival time predictions.

## MATERIAL & DATA AVAILABILITY

All data, including the 1013 patient cohort, relevant scripts and PDACNet is available at: https://github.com/KristoferLintonReid/Enhanced-Cancer-Subtyping-

## ACKNOWLEDGMENTS

The authors would like to thank everyone at and involved with Cambridge Cancer Genomics whom made this project possible and offered insightful constructive feedback throughout this study.

## APPENDIX

**Figure 1.**
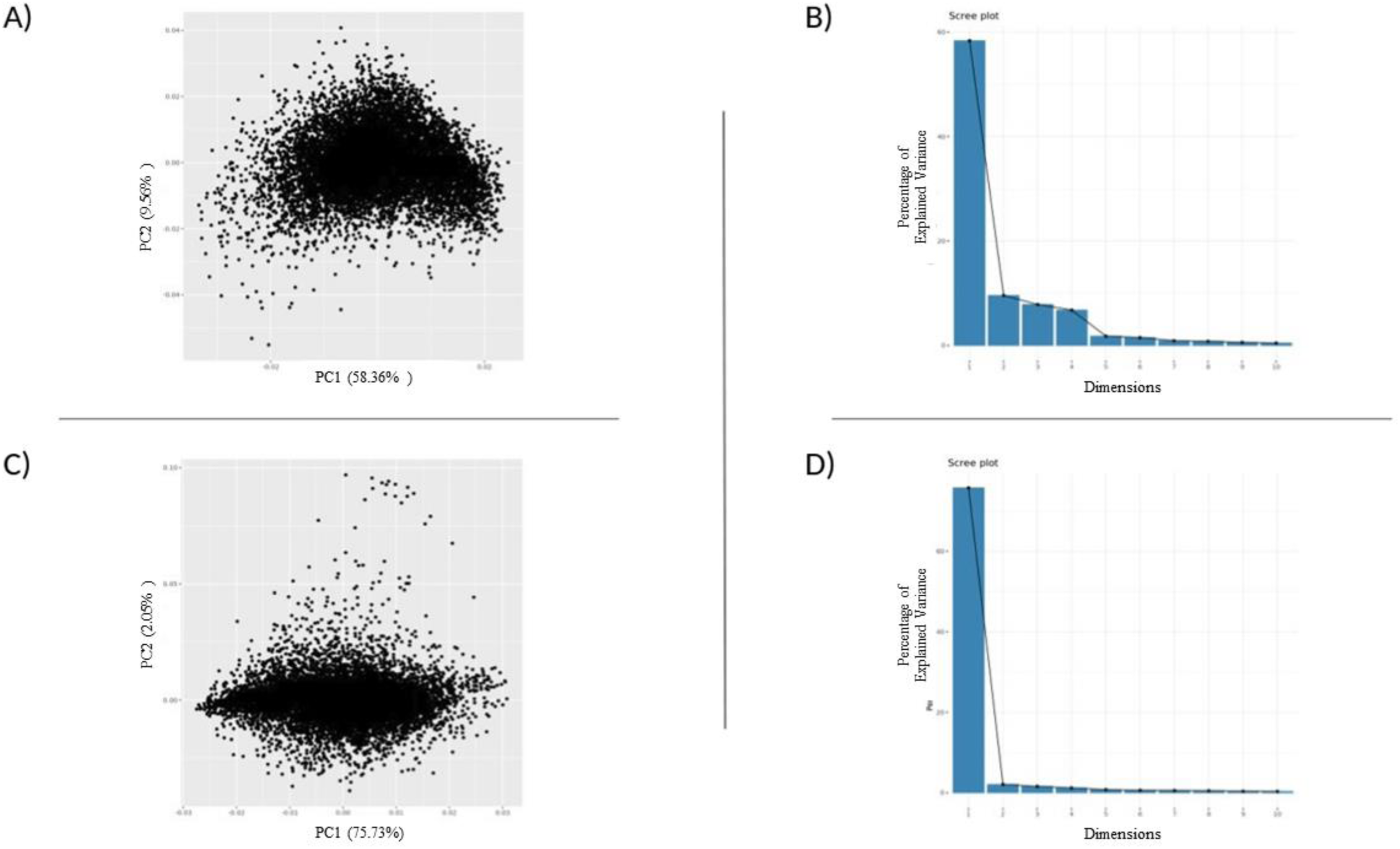
Combined RNA-Sequencing Data Score and Scree Plots (Components 1 &2). A) Score plot before ComBat bath correction. B) Scree Plot Pre-ComBat batch correction. C) Score Plot of ComBat corrected RNA-Seq Data. D) Scree Plot of ComBat corrected data.

**Figure 2.**
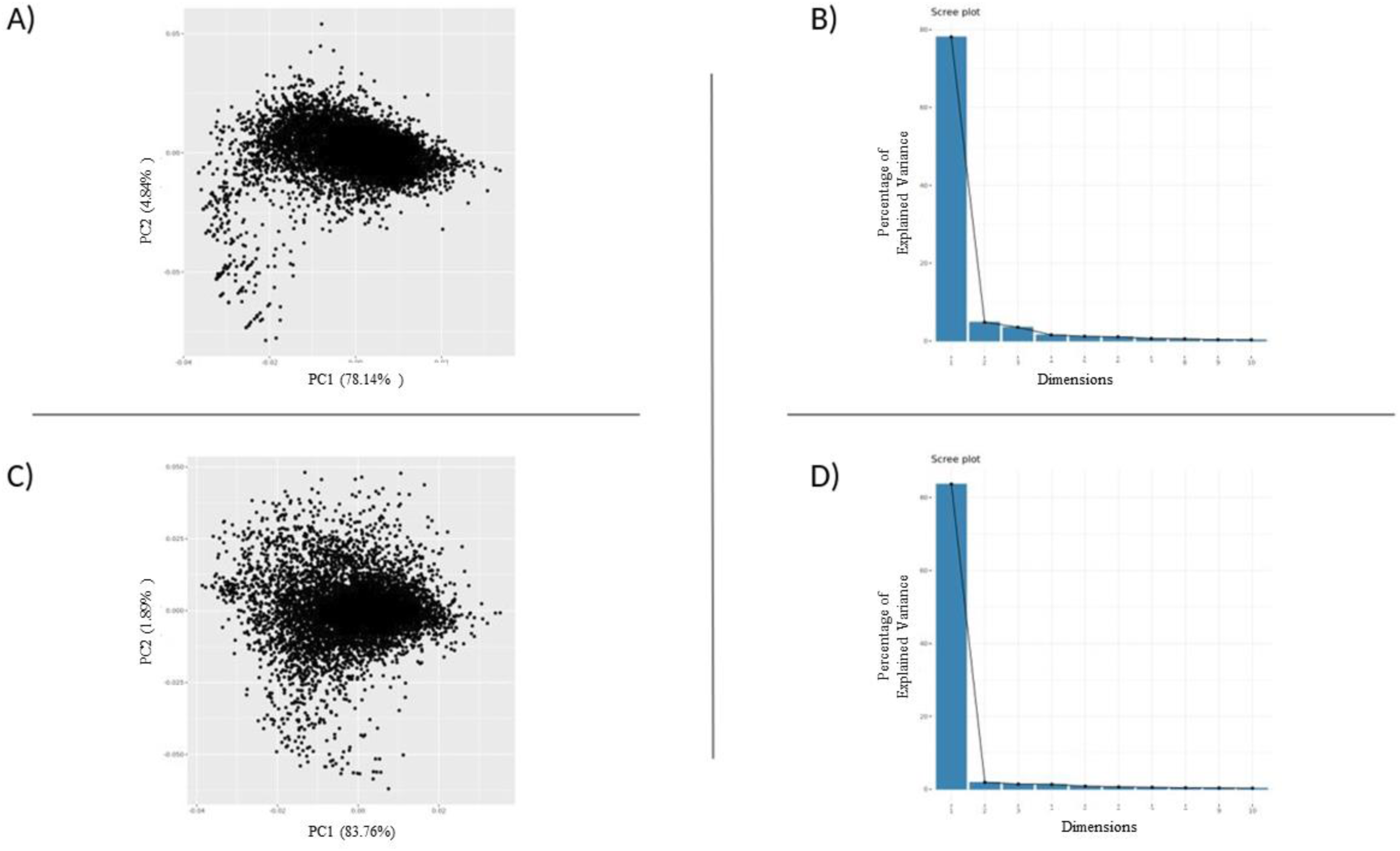
Combined Microarray Data Score and Scree Plots (Components 1 &2). A) Score plot before ComBat bath correction. B) Scree Plot Pre-ComBat batch correction. C) Score Plot of ComBat corrected Microarray Data. D) Scree Plot of ComBat corrected data.

**Figure 3.**
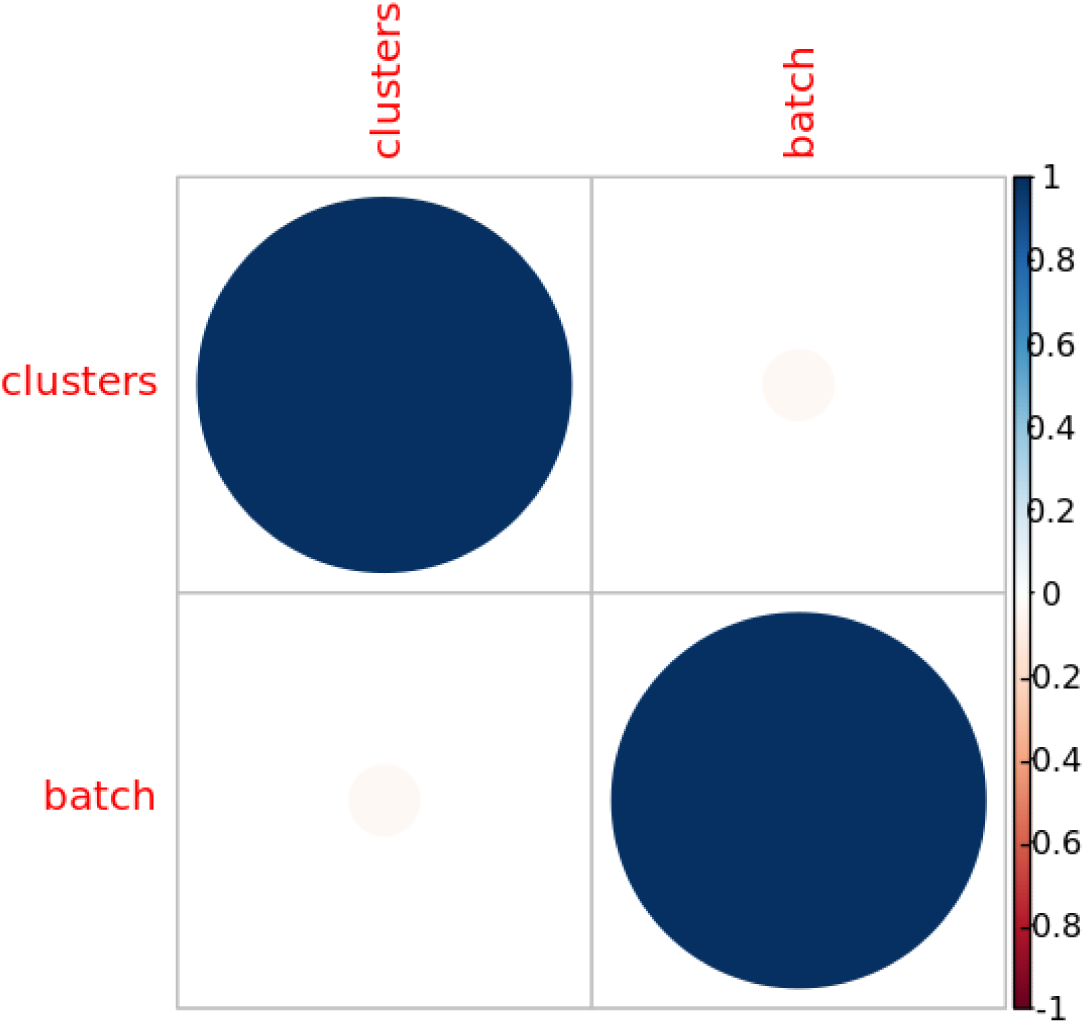
Pearson’s Correlation between batches and M3C distinguished clusters. t= −1.1188, p= 0.2635, cor= - .003535712. There is slight negative correlation, however not significant.

**Table 1.** Top 5 Significantly Differentiated Genes of Each Subtype.

|  |  | logFC | AveExpr (log) | t | P.Value | adj.P.Val | B |
| --- | --- | --- | --- | --- | --- | --- | --- |
| CLASS 1 | AOX1 | -0.2697102765 | 7.383402014 | -24.71732052 | 1.05E-105 | 6.97E-102 | 230.3889938 |
|  | FLRT2 | -0.2343510896 | 7.422330274 | -23.15971162 | 2.18E-95 | 7.27E-92 | 206.7464655 |
|  | SLC1A2 | -0.2736117953 | 7.197830691 | -22.22002319 | 2.94E-89 | 5.22E-86 | 192.7045101 |
|  | GAMT | -0.1698139592 | 7.439118734 | -22.21577758 | 3.13E-89 | 5.22E-86 | 192.6414979 |
|  | KCNJ5 | -0.2170978977 | 7.268332395 | -21.5335573 | 7.79E-85 | 9.93E-82 | 182.5707715 |
| CLASS 2 | GAMT | 0.139585 | 7.439119 | 21.22054 | 7.79E-83 | 5.20E-79 | 177.955 |
|  | P2RX1 | 0.200646 | 7.318828 | 20.76559 | 6.02E-80 | 2.01E-76 | 171.3457 |
|  | NUCB2 | 0.134035 | 7.564929 | 19.83386 | 4.11E-74 | 9.14E-71 | 157.9919 |
|  | RBPJL | 0.240064 | 7.297063 | 19.44479 | 1.04E-71 | 1.73E-68 | 152.4936 |
|  | S100A6 | -0.11722 | 7.674083 | -19.4097 | 1.71E-71 | 2.28E-68 | 152.0004 |
|  |  | logFC*(-) |  |  |  |  |  |
| CLASS 3 | GAMT | -0.13959 | 7.439119 | -21.2205 | 7.79E-83 | 5.20E-79 | 177.955 |
|  | P2RX1 | -0.20065 | 7.318828 | -20.7656 | 6.02E-80 | 2.01E-76 | 171.3457 |
|  | NUCB2 | -0.13403 | 7.564929 | -19.8339 | 4.11E-74 | 9.14E-71 | 157.9919 |
|  | RBPJL | -0.24006 | 7.297063 | -19.4448 | 1.04E-71 | 1.73E-68 | 152.4936 |
|  | S100A6 | 0.117216 | 7.674083 | 19.40973 | 1.71E-71 | 2.28E-68 | 152.0004 |
| CLASS 4 | EPB41L4B | -0.19721 | 7.449825 | -19.9049 | 1.49E-74 | 9.93E-71 | 158.9589 |
|  | RAB31 | 0.120711 | 7.595502 | 19.19366 | 3.60E-70 | 1.20E-66 | 148.9385 |
|  | CDH11 | 0.14974 | 7.544486 | 18.75932 | 1.57E-67 | 3.50E-64 | 142.9014 |
|  | NUCB2 | -0.13217 | 7.564929 | -17.9862 | 6.68E-63 | 1.11E-59 | 132.3218 |
|  | RBPJL | -0.23903 | 7.297063 | -17.8049 | 7.87E-62 | 1.05E-58 | 129.8736 |
| CLASS 5 | CPA2 | -0.38955 | 7.48819 | -22.2839 | 1.13E-89 | 7.55E-86 | 193.6168 |
|  | PLA2G1B | -0.3834 | 7.50473 | -22.0594 | 3.22E-88 | 1.07E-84 | 190.2892 |
|  | CTRB2 | -0.30308 | 7.595823 | -22.007 | 7.01E-88 | 1.56E-84 | 189.5151 |
|  | CEL | -0.38869 | 7.446912 | -21.8277 | 1.00E-86 | 1.68E-83 | 186.8672 |
|  | CUZD1 | -0.34999 | 7.43262 | -21.3858 | 6.87E-84 | 9.16E-81 | 180.3775 |
| CLASS 6 | AP1M2 | -0.1625 | 7.441295 | -20.5203 | 2.12E-78 | 1.41E-74 | 167.7724 |
|  | EPCAM | -0.15343 | 7.670752 | -20.1773 | 2.99E-76 | 9.99E-73 | 162.8536 |
|  | ATP1B1 | -0.1353 | 7.716065 | -19.1836 | 4.15E-70 | 9.21E-67 | 148.8091 |
|  | PLS1 | -0.20998 | 7.52294 | -19.0886 | 1.58E-69 | 2.63E-66 | 147.4828 |
|  | CDS1 | -0.19613 | 7.441095 | -18.9602 | 9.53E-69 | 1.27E-65 | 145.6956 |

**Table 2.** Performance of the Top Classification Model. A total error of 0.186047, at a rate of 56/301.

|  | Class1 | Class2 | Class3 | Class4 | Class5 | Class6 | Error | Rate |
| --- | --- | --- | --- | --- | --- | --- | --- | --- |
| Class1 | 36 | 0 | 3 | 0 | 8 | 1 | 0.25 | 12/48 |
| Class2 | 0 | 20 | 5 | 1 | 0 | 0 | 0.230769 | 6/26 |
| Class3 | 2 | 3 | 87 | 7 | 4 | 0 | 0.15534 | 16 / 103 |
| Class4 | 0 | 0 | 7 | 45 | 2 | 4 | 0.224138 | 13 / 58 |
| Class5 | 1 | 0 | 2 | 1 | 39 | 0 | 0.093023 | 4/43 |
| Class6 | 0 | 0 | 0 | 4 | 1 | 18 | 0.217391 | 5/23 |
| Totals | 39 | 23 | 104 | 58 | 54 | 23 | 0.186047 | 56 / 301 |

**Table 3.** Proportion of batches in each subtype.

|  | Proportion of<br>Batches in<br>Class 1 (%) | Proportion of<br>Batches in<br>Class 2 (%) | Proportion of<br>Batches in<br>Class 3 (%) | Proportion of<br>Batches in<br>Class 4 (%) | Proportion of<br>Batches in<br>Class 5 (%) | Proportion of<br>Batches in<br>Class 6 (%) |
| --- | --- | --- | --- | --- | --- | --- |
| Batch 1 | 0 | 1 | 5 | 4 | 2 | 0 |
| Batch 2 | 2 | 0 | 9 | 9 | 0 | 0 |
| Batch 3 | 2 | 6 | 3 | 2 | 3 | 12 |
| Batch 4 | 5 | 9 | 4 | 5 | 5 | 5 |
| Batch 5 | 12 | 13 | 12 | 12 | 13 | 8 |
| Batch 6 | 4 | 11 | 6 | 2 | 12 | 12 |
| Batch 7 | 20 | 16 | 12 | 12 | 15 | 17 |
| Batch 8 | 5 | 9 | 9 | 7 | 10 | 5 |
| Batch 9 | 6 | 5 | 5 | 8 | 5 | 3 |
| Batch 10 | 5 | 5 | 4 | 2 | 7 | 7 |
| Batch 11 | 5 | 4 | 2 | 5 | 5 | 1 |
| Batch 12 | 7 | 6 | 9 | 3 | 7 | 8 |
| Batch 13 | 21 | 16 | 16 | 19 | 16 | 17 |
| Batch 14 | 8 | 2 | 5 | 10 | 1 | 4 |

**Figure 4:**
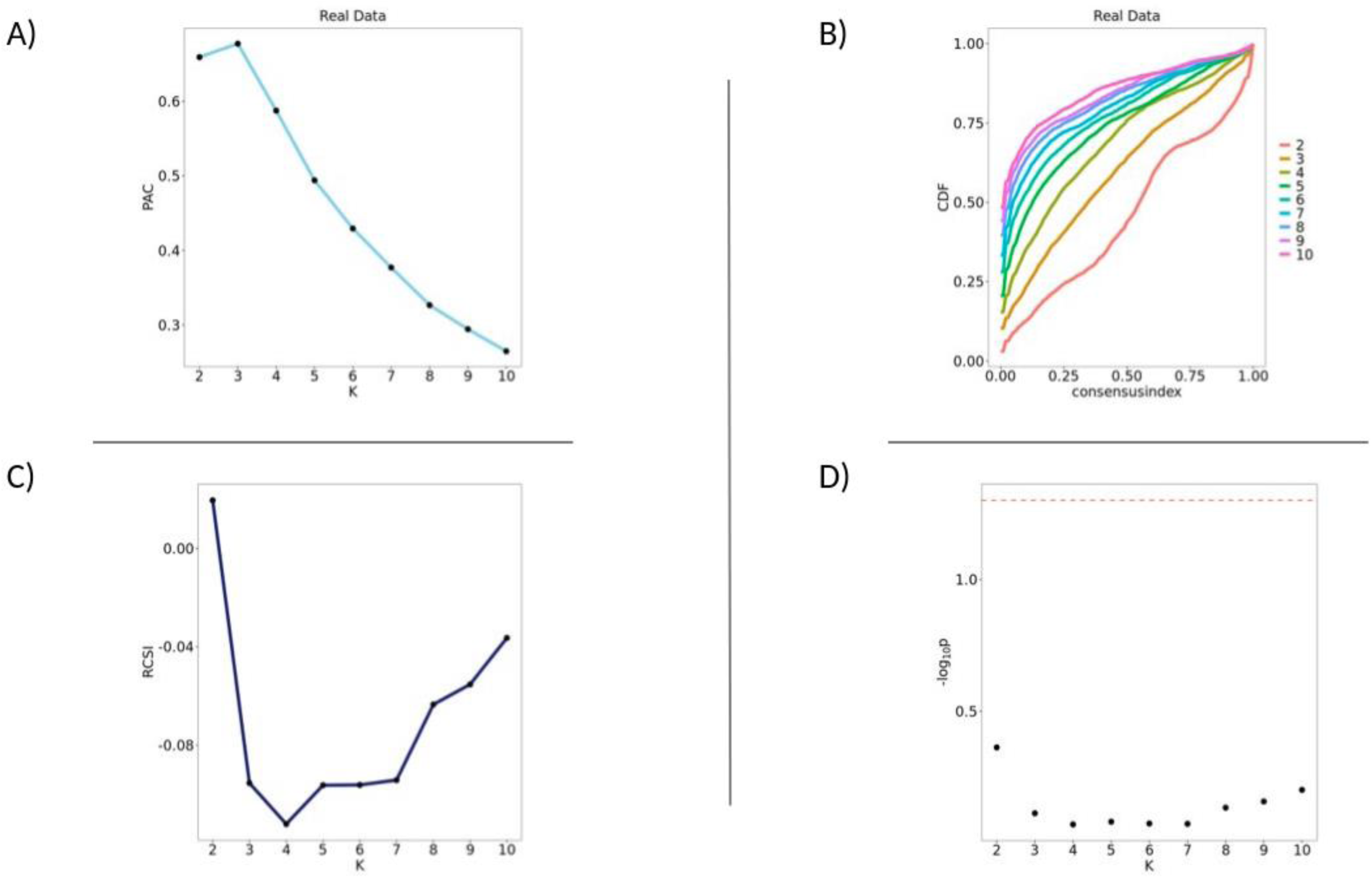
M3C algorithm applied to the cohort form Bailey et al. [1]. A) PAC scores display a sharp upwards spike at K=3, and a slight arch at K=4. B) CDF plot shows flattening of the consensus index curve as K increases. C) RSCI shows a sharp downwards spike at K=4. D)Plot of the P-value distribution over a logical distribution of K values. Here, none of the ranks are significant.

**Figure 5:**
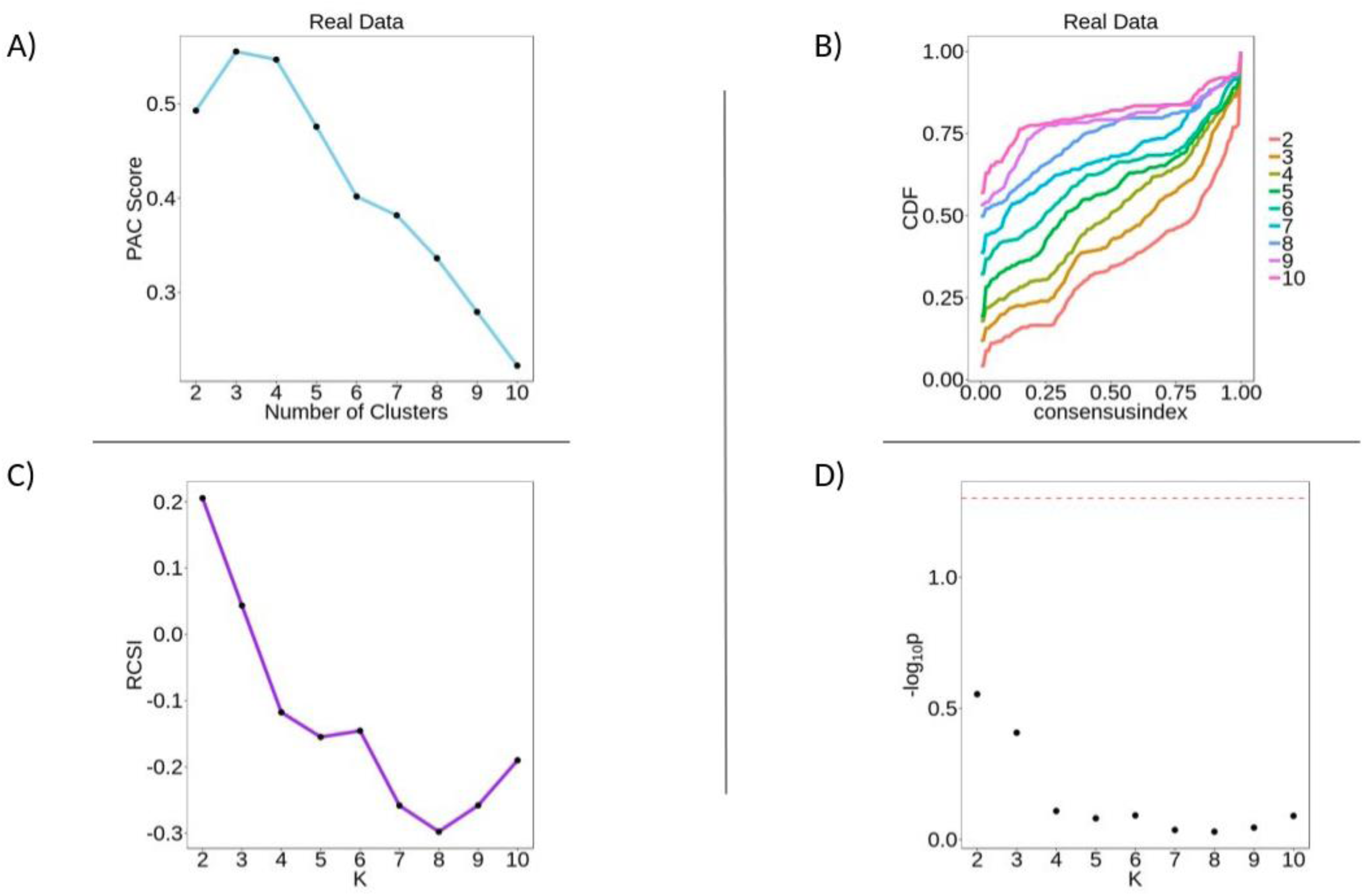
Collisson’s Cohort M3C Results. M3C analysis applied to the cohort from Collisson et al., [2]. A) PAC scores display a sharp upwards spike at K=3, and a slight arch at K=4. B) CDF plot shows flattening of the consensus index curve at K increases. C) RSCI shows a sharp downwards spike at K= 4 and 7, with arching between K=5:8. D) Plot of the P-values over a logical distribution of K values. None of the ranks were above the significance threshold.

